# Rampant tooth loss across 200 million years of frog evolution

**DOI:** 10.1101/2021.02.04.429809

**Authors:** Daniel J. Paluh, Karina Riddell, Catherine M. Early, Maggie M. Hantak, Gregory F.M. Jongsma, Rachel M. Keeffe, Fernanda Magalhães Silva, Stuart V. Nielsen, María Camila Vallejo-Pareja, Edward L. Stanley, David C. Blackburn

## Abstract

Teeth have been broadly maintained across most clades of vertebrates but have been lost completely at least once in actinopterygian fishes and several times in amniotes. Using phenotypic data collected from over 500 genera via micro-computed tomography, we provide the first rigorous assessment of the evolutionary history of dentition across all major lineages of amphibians. We demonstrate that dentition is invariably present in caecilians and salamanders, but teeth have been lost completely more than 20 times in frogs, a much higher occurrence of edentulism than in any other vertebrate group. The repeated loss of teeth in anurans is associated with a specialized diet of small invertebrate prey as well as shortening of the lower jaw, but it is not correlated with a reduction in body size. Frogs provide an unparalleled opportunity for investigating the molecular and developmental mechanisms of convergent tooth loss on a large phylogenetic scale.

## Introduction

The evolution of teeth is considered a key innovation that promoted the radiation of jawed vertebrates, facilitating the transition from a passive to active predatory lifestyle (Gans and Northcutt 1983). Teeth are complex mineralized tissues that originated in stem gnathostomes more than 400 million years ago (Rücklin et al. 2012) and have been broadly maintained across living chondrichthyans, actinopterygians, and sarcopterygians due to the critical role these structures play in the acquisition and processing of food. The shape, size, location, and number of teeth differ widely across vertebrates, especially in response to broad variation in food type. Although dentition is generally conserved across vertebrates, teeth have been lost completely several times, resulting in toothlessness or edentulism, including in three extant clades of mammals (baleen whales, anteaters, and pangolins), turtles, and birds (Davit-Béal et al. 2009). Teeth are likely lost following the evolution of a secondary feeding tool that improves the efficiency of food intake (e.g., beak, baleen, specialized tongue), leading to relaxed functional constraints on dentition (Davit-Béal et al. 2009). In contrast to other tetrapods, the evolution and diversity of teeth in amphibians has been poorly studied, despite long recognition that frogs— one of the most diverse vertebrate orders with more than 7,000 species—possess variation in the presence or absence of teeth.

All living salamanders and caecilians are assumed to have teeth on the upper jaw, lower jaw, and palate (Duellman and Trueb 1986), but nearly all frogs lack dentition on the lower jaw and variably possess teeth on the upper jaw and palate. Recent work suggests that dentition on the lower jaw was lost in the ancestor of frogs more than 200 million years ago and was subsequently regained in a single species (*Gastrotheca guentheri*; Boulenger 1882) during the Miocene (Wiens 2011). The presence or absence of dentition has also been considered an important taxonomic character in frogs; for example, a subclass was once proposed that included all toothless species (Bufoniformia; Cope 1867). Our understanding of the anuran tree of life has fundamentally changed with the development of molecular phylogenetics (Duellman and Trueb 1986, Feng et al. 2017; Hime et al. 2020), but there has been no attempt to estimate the frequency of tooth loss across frog diversity or evaluate the factors that may be correlated with edentulism. Most frogs are generalist, gape-limited predators that capture prey using tongue propulsion (Regal and Gans 1976), reducing the importance of teeth in prey capture. Tooth loss is hypothesized to occur in frogs that specialize on eating small prey (microphagy), such as ants and termites (Das and Coe 1994, Parmelee 1999, Narvaez and Ron 2013). This may lead to relaxed functional constraints on energetically expensive teeth. Microphagous frogs are known to have shortened jaws and altered feeding cycles (Emerson 1985), modified tongues (Trueb and Gans 1983), and some have the ability to sequester dietary alkaloids from their prey, rendering them toxic (Caldwell 1996, Vences et al. 1998). Alternately, teeth may be reduced or lost as a byproduct of miniaturization or truncated development (paedomorphosis; Davies 1989, Hanken and Wake 1993, Smirnov and Vasil’eva 1995) because the initiation of odontogenesis occurs ontogenetically late in frogs (during or after metamorphosis) compared to other vertebrates.

Using the most recent species-rich phylogeny of extant amphibian species (Jetz and Pyron 2018) and our extensive taxonomic sampling via high-resolution X-ray micro-computed tomography of over 500 of the 561 currently recognized amphibian genera (AmphibiaWeb 2021), we 1) evaluated the phylogenetic distribution of teeth and reconstructed the evolutionary history of dentition across all major lineages of amphibians and 2) tested whether dietary specialization, relative jaw length, and body size are correlated with the loss of teeth in frogs. Our results demonstrate that the presence and location of teeth are highly conserved in salamanders and caecilians, but labile in frogs. We found that teeth have been repeatedly lost in frogs and at a much higher frequency than in any other vertebrate group. The evolution of edentulism in anurans is correlated with a microphagous diet and shortening of the lower jaw but not with a reduction in body size over evolutionary time. Six reversals, from edentulous to toothed jaws, were inferred in frogs.

## Results

### Distribution of teeth in amphibians

We recorded the presence or absence of teeth on each dentigerous bone of the lower jaw, upper jaw, and palate for 524 amphibian species (Fig. 1; Dataset S1). Taxa were coded as “toothed” if teeth were observed on any cranial element and “edentulous” if teeth were entirely absent. Our survey of amphibian dentition across the majority of extant genera confirmed that all salamanders and caecilians retain teeth, while 134 of the 429 frog species examined are entirely edentulous (Dataset S1). All anuran species lack dentary teeth with the exception of *Gastrotheca guentheri*. Maxillary and premaxillary teeth of the upper jaw co-occur in all frog species (Fig. 1A, 1B), being present in 292 taxa and absent in 136 species. The vomerine teeth on the palate are the most variable in frogs, being present in 202 species and absent in 226. Many anurans have maxillary and premaxillary teeth in the absence of vomerine teeth (92 species), but only two species examined have vomerine teeth while lacking upper jaw teeth (*Rhombophryne testudo, Uperodon systoma*).

**Figure 1.**
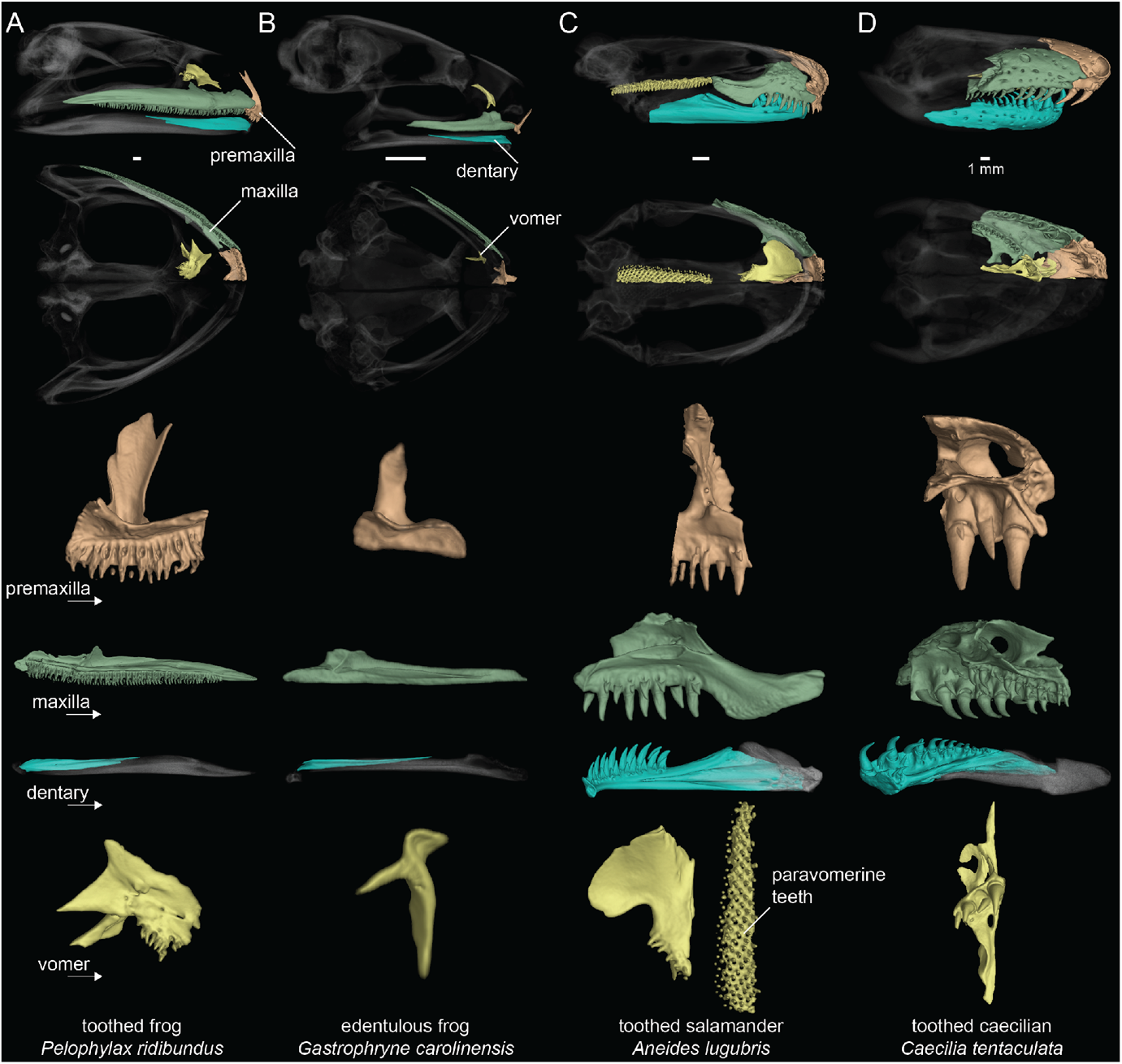
Dental diversity of amphibians. **A**. Toothed frog, *Pelophylax ridibundus* (CAS:Herp:217695), **B**. edentulous frog, *Gastrophryne carolinensis* (UF:Herp:110645), **C**. toothed salamander, *Aneides lugubris* (MVZ:Herp:249828), **D**. toothed caecilian, *Caecilia tentaculata* (KU:Kuh:175441). Skulls in lateral and ventral views: dentigerous cranial elements are colored and the remainder of the skull is semi-transparent. Isolated premaxilla (orange), maxilla (green), and dentary (blue) in lingual views. Isolated vomer (yellow) in ventral views. Teeth are present on all colored elements except the dentary in *P. ridibundus* and those of *G. carolinensis*. Scale bars = 1 mm.

All 65 salamander species examined have teeth on the lower jaw and palate, but three species lack upper jaw teeth on the maxilla and premaxilla (the sirenids *Siren intermedia* and *Pseudobranchus striatus* and the salamandrid *Salamandrina terdigitata*). *Thorius pennatulus* (Plethodonidae) and two proteids (*Necturus lewisi* and *Proteus anguinus*) lack maxillary teeth but retain premaxillary teeth. All salamanders have vomerine teeth on the palate (including the paravomerine tooth patches that underlie the parasphenoid in plethodontids (Fig. 1C); Lawson et al. 1971). Palatal teeth were additionally observed on the palatopterygoid (*Necturus lewisi* and *Proteus anguinus*) and palatine (*Siren intermedia* and *Pseudobranchus striatus*). The lower jaw teeth are present on the dentary in all species, except *Siren intermedia* and *Pseudobranchus striatus*, which have mandibular teeth on the splenial. *Necturus lewisi* and *Proteus anguinus* are the only two species that have lower jaw teeth on both the dentary and splenial.

All 30 caecilian species examined possess teeth on the lower jaw, upper jaw, and palate (Fig. 1D). The individual elements of the lower jaw in caecilians fuse to form the pseudodentary, and this composite element varies in having either one or two rows of teeth. Upper jaw teeth are present on the nasopremaxilla (fused nasal and premaxilla) and maxillopalatine (fused maxilla and palatine; outer row). Palatal teeth are always present on the vomer and maxillopalatine (inner row) and occur on the ectopterygoid in one species (*Geotrypetes seraphini*).

### Repeated Tooth Loss in Frogs

Teeth are absent in 134 anuran genera belonging to 19 families. We used reversible-jump Markov chain Monte Carlo (MCMC) in RevBayes (Höhna et al. 2016) to sample all five Markov models of phenotypic character evolution in proportion to their posterior probability. The maximum a posteriori model of dentition evolution was the one-rate model with a posterior probability of 0.91. The model-averaged maximum a posteriori ancestral state of Lissamphibia and Anura is toothed with a posterior probability of 0.99. Teeth have been completely lost at least 22 times in frogs (Fig. 2), and six reversals from edentulous to toothed upper jaws were inferred. Edentulism has evolved three times in Mesobatrachia, 12 times in Hyloidea, six times in Ranoidea, and once in Nasikabatrachidae. One reversal was estimated in Myobatrachidae (in *Uperoleia mahonyi*; Clulow et al. 2016) and five reversals were inferred in Microhylidae (in *Dyscophus, Uperodon, Anodonthyla, Cophyla*, and *Rhombophryne* + *Plethodontohyla*).

**Figure 2.**
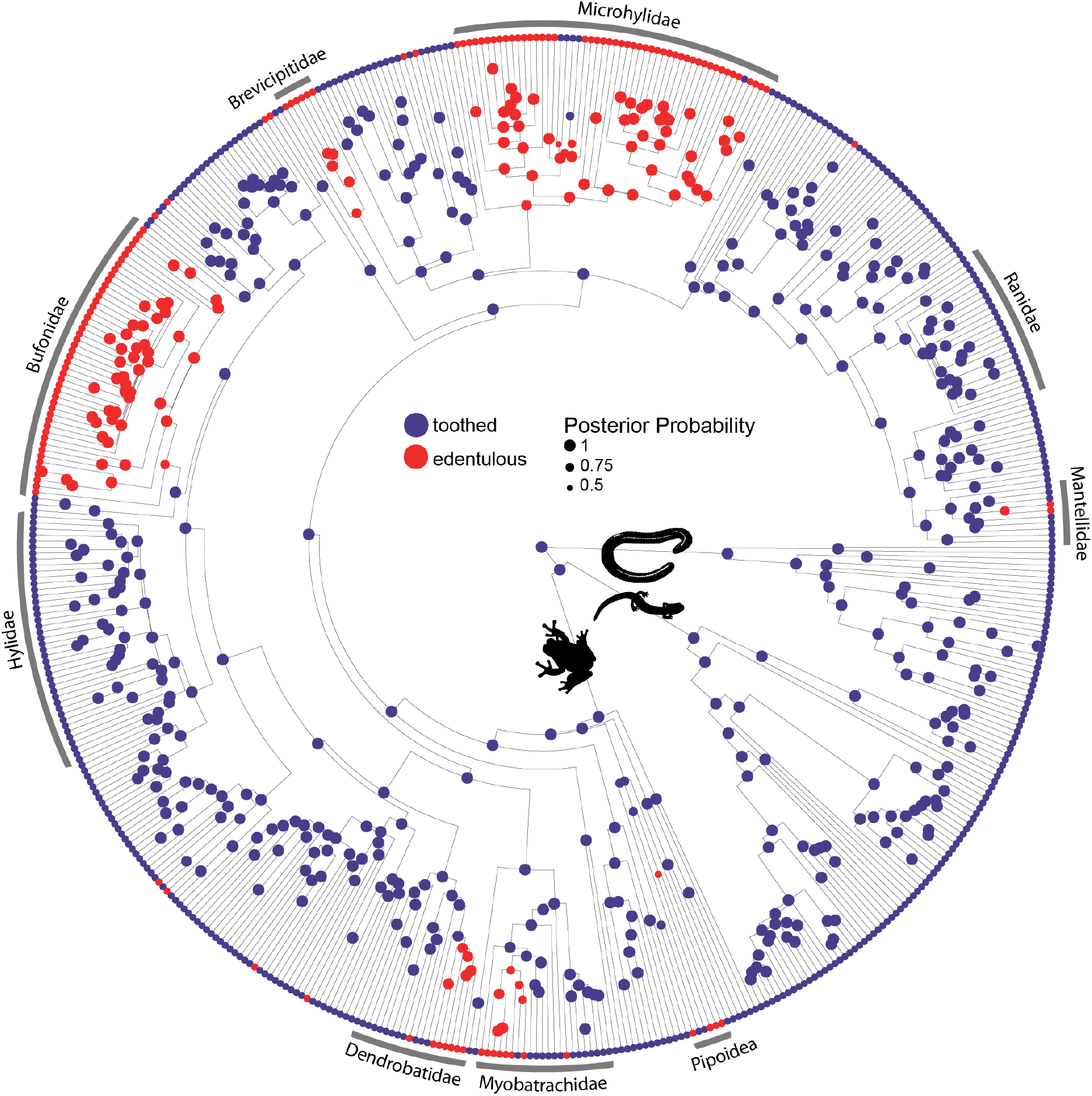
Phylogeny of 524 amphibians depicting the evolution of dentition. Node point color corresponds to Bayesian model-averaged ancestral states of dentition: blue = toothed; red = edentulous. The size of each node point represents the posterior probability of the most probable ancestral state. Tip point colors correspond to dentition states for all species. For species tip labels display Figure 1—figure supplement 1. For Bayesian model-averaged ancestral states of tooth presence/absence on individual dentigerous elements display Figure 1—figure supplements 2–5. Corresponding data are provided in Dataset S1.

### Relationships among tooth loss, diet, and body size

We compiled published diet records for 267 frog lineages and classified 69 taxa from 20 families as microphagous and 198 taxa from 47 families as generalist feeders (Fig. 3; Dataset S2). Of the 69 microphagy specialists, 53 are edentulous and 16 are toothed. Of the 198 generalists, 26 are edentulous and 172 are toothed. A BayesTrait discrete analysis indicated correlated evolution between edentulism and microphagy: the dependent model of trait evolution is strongly supported over the independent model (Bayes factor = 46.97; a Bayes factor > 2 implies the evolution of two traits is linked). Similar results were found using a 158-taxon dataset excluding genus-level diet data (Bayes factor = 26.26). Of the 22 independent losses of teeth across frogs, at least 16 of these lineages contain microphagous species (Fig. 3). The majority of the 26 taxa classified as both edentulous and generalist feeders are members of the Bufonidae and Microhylidae, but also includes the fully aquatic pipids *Pipa* and *Hymenochirus*, two brevicipitids (*Probreviceps* and *Callulina*), and the Darwin’s frog, *Rhinoderma*.

**Figure 3.**
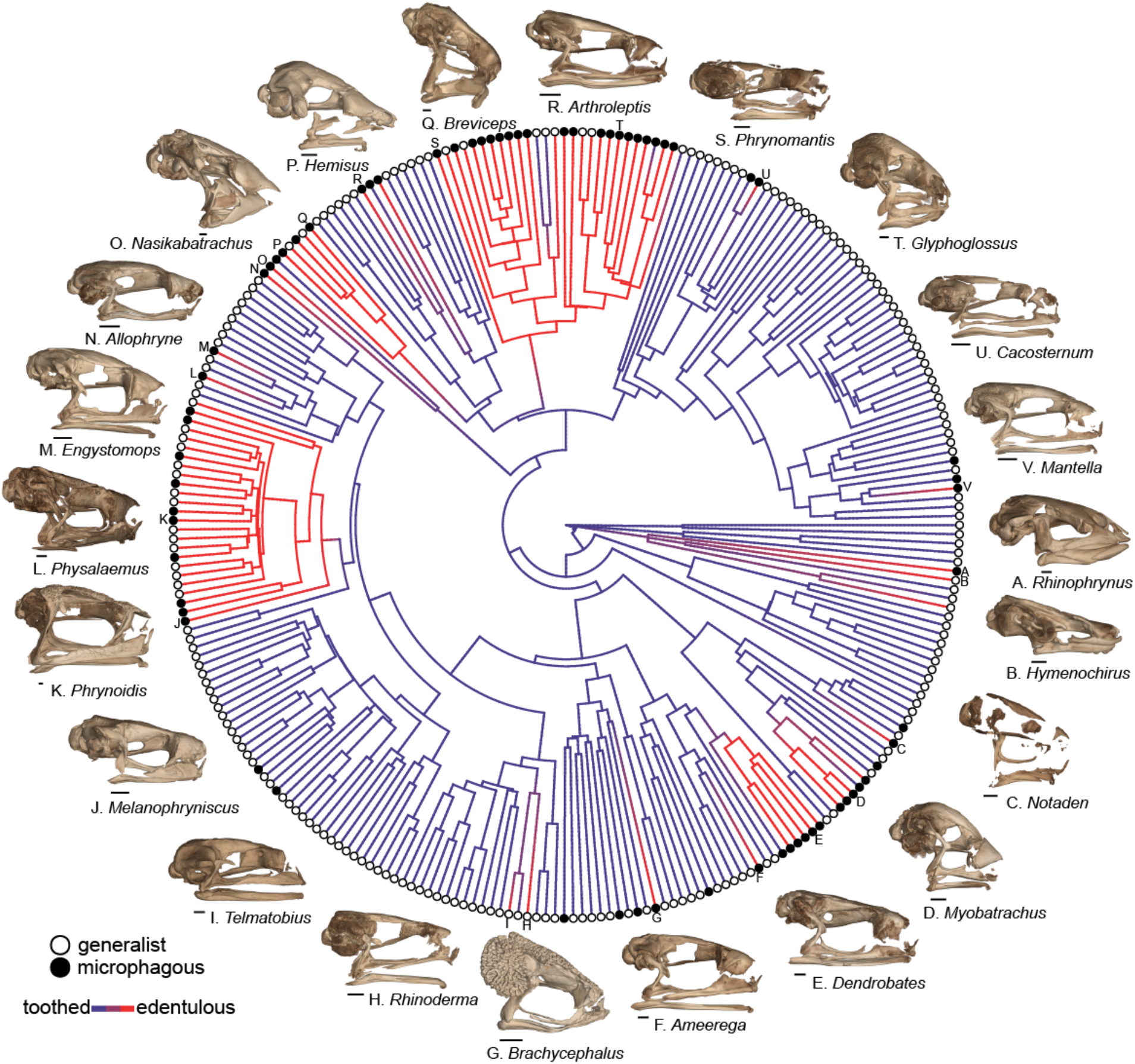
Phylogeny of 267 frog species with a stochastic character map of dentition states (phytools, Revell 2012) and distribution of generalist and microphagous diet states (tip point colors) illustrating the correlated evolution of edentulism and microphagy. Diversity of edentulous frog skulls: **A**. *Rhinophrynus dorsalis* (CAS:Herp:71766), **B**. *Hymenochirus boettgeri* (CAS:Herp:253587), **C**. Notaden bennetti (CAS:Herp:78115), **D**. Myobatrachus gouldii (MCZ:Herp:139543), **E**. *Dendrobates tinctorius* (YPM:Vz:010610), **F**. *Ameerega trivittata* (UF:Herp:107200), **G**. *Brachycephalus ephippium* (UF:Herp:72725), **H**. *Rhinoderma darwinii* (UF:Herp:62022), **I**. *Telmatobius carrillae* (UF:Herp:39717), **J**. *Melanophryniscus stelzneri* (UF:Herp:63183), **K**. *Phrynoidis asper* (USNM:Amphibians & Reptiles:586870), **L**. *Physalaemus nattereri* (MCZ:Herp:A30113), **M**. *Engystomops pustulosus* (CAS:Sua:21892), **N**. *Allophryne ruthveni* (KU:Kuh:166716), **O**. *Nasikabatrachus sahadryensis* (CES:F:203), **P**. *Hemisus guineensis* (CAS:Herp:258533), **Q**. *Breviceps gibbosus* (AMNH:Herpetology:3053), **R**. *Arthroleptis schubotzi* (CAS:Herp:201762), **S**. *Phrynomantis annectens* (AMB:10086), **T**. *Glyphoglossus molossus* (CAS:Herp:243121), **U**. *Cacosternum namaquense* (CAS:Herp:156975), **V**. *Mantella baroni* (CAS:Herp:250387). Scale bars = 1 mm. For species tip labels display Figure 2—figure supplement 1. Corresponding data are provided in Dataset S2.

The relative jaw length in frogs ranges from 62% of head length in *Synapturanus mirandaribeiroi*, an edentulous microhylid, to 140% of head length in *Lepidobatrachus asper*, a toothed ceratophryid. A phylogenetic logistic regression showed a significant relationship between edentulism and shortened jaws (alpha = 0.0011, standard error = 0.8920, *P* < 0.001; Fig. 4A). Edentulous species have an average relative jaw length of 83% of head length, while toothed species have an average relative jaw length of 99% of head length. Nearly all edentulous species examined have an anteriorly shifted jaw joint (lower jaw length is shorter than respective head length; the two largest bufonids in our dataset, *Bufo gargarizans* and *Rhaebo blombergi*, are exceptions), but over 100 toothed taxa have posteriorly shifted jaws (lower jaws that are longer than their heads; Fig. 4A). The snout–vent length (SVL) of specimens measured ranges from 7.8 mm in the edentulous *Paedophryne amauensis*, the smallest known frog, to 263.9 mm in the toothed *Conraua goliath*, the largest known extant species. A phylogenetic logistic regression indicated that there is no relationship between edentulism and body size (alpha = 0.0016, standard error = 0.1162, *P* = 0.09; Fig 4B). Edentulous species have an average SVL of 36.2 mm (range 7.8–152.4 mm) and toothed species have an average SVL of 43.7 (range 11.2–263.9 mm).

**Figure 4.**
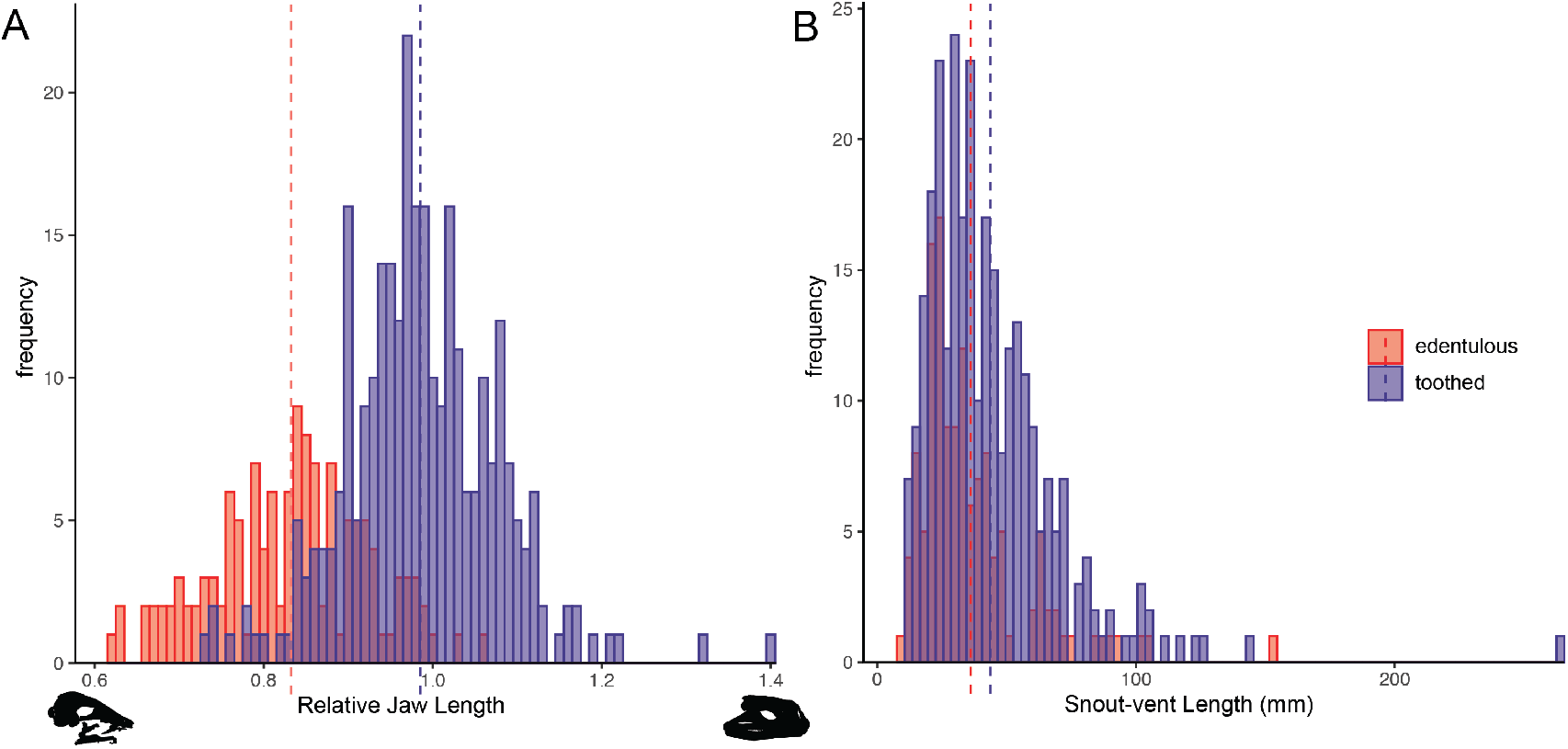
Histograms of relative jaw length (mandible length divided by skull length; **A**) and body size (snout-vent length; **B**) in 423 frog species plotted by dentition states (blue = toothed; red = edentulous). A phylogenetic correlation was identified between tooth loss and shortened lower jaws. There is no association between edentulism and body size. Left skull silhouette is *Hemisus guineensis* (CAS:Herp:258533) and right skull is *Lepidobatrachus asper* (UF:Herp:12347). Corresponding trait data are provided in Dataset S3.

## Discussion

### Evolution of edentulism in jawed vertebrates

With at least 22 independent origins of edentulism, frogs have completely lost teeth more times than any other vertebrate clade. Based on our review of the literature, only seven other extant vertebrate lineages are entirely edentulous. There are no described edentulous chondrichthyan species (but see Mulas et al. (2020) for the first described aberrant case in a catshark). To our knowledge, teeth have been entirely lost only twice in living actinopterygian fishes in the Syngnathidae (seahorses and pipefish; Lin et al. 2016) and the milkfish, *Chanos chanos* (Kohno et al. 1996, Wang et al. 2017). Other fish lineages, such as the cyprinids, have toothless oral jaws but retain true pharyngeal teeth (Aigler et al. 2014). Five extant amniote clades are edentulous, including three lineages of mammals (baleen whales, pangolins, anteaters), all living birds, and all living turtles (Davit-Béal et al. 2009). There are several mammal clades that have lost enamel but retain reduced teeth (armadillos, sloths, aardvarks, pygmy and dwarf sperm whales; Meredith et al. 2009). Molecular evidence suggests a single loss of teeth in the common ancestor of extant birds (Meredith et al. 2014), but complete edentulism also evolved independently in at least two extinct lineages of Mesozoic birds (*Confuciusornis* and *Gobipteryx*; Yang and Snyder 2018). Teeth were completely lost in at least two lineages of non-avian dinosaurs (ornithomimosaurs and caenagnathoids; Wang et al. 2017, Hendrickx et al. 2019) and in some pterosaurs, such as members of Azhdarchidae (Yang and Snyder 2018). All living crocodilians retain teeth, but at least two fossil suchian archosaurs were edentulous (*Shuvosaurus* and *Effigia*; Nesbitt and Norell 2006). There are no known edentulous squamate species, although African egg-eating snakes in the genus *Dasypeltis* may have a dental polymorphism, as they typically have small, short teeth but some individuals are reported to be edentulous (Visser 1981). Lastly, at least one extinct rhynchocephalian has been suggested to be edentulous (*Sapheosaurus*, Rauhut et al. 2012).

The loss of teeth may be associated with the evolution of a secondary feeding apparatus (Davit-Béal et al. 2009, Wang et al. 2017), such as the keratinized beak in birds and turtles, baleen in mysticete whales, and specialized tongues in pangolins and anteaters. Nearly all frogs have a specialized tongue that is used in feeding (Regal and Gans 1976), and this adaptation might have facilitated the repeated loss of teeth across anurans. Surprisingly, three anuran lineages are both tongueless and edentulous (*Hymenochirus, Pseudhymenochirus*, and *Pipa* in Pipidae), but these species are highly aquatic and have a derived mechanism of catching prey under water through suction feeding (Dean 2003, Cundall et al. 2017). The edentulous syngnathids (seahorses and relatives) and actinopterygians with toothless oral jaws also catch prey through suction feeding (Roos et al. 2009, Mihalitsis and Bellwood 2019). There appears to be no size-related constraints promoting complete tooth loss across all vertebrates. Edentulous species span the entire spectrum of vertebrate body sizes: the smallest known vertebrate species (the microhylid frog *Paedophryne amauensis*, Rittmeyer et al. 2012) and the largest (the blue whale, *Balaenoptera musculus*) are both edentulous. The second-smallest known vertebrate, the cyprinid fish in the genus *Paedocypris*, retain true pharyngeal teeth (Kottelat et al. 2006). Several edentulous vertebrate clades are thought to have paedomorphic skulls, including toothless frogs (Smirnov and Vasil’eva 1995), birds (Bhullar et al. 2012), and baleen whales (Fordyce and Barnes 1994), suggesting that tooth loss in vertebrates may be a byproduct of truncated development, but this hypothesis requires further investigation.

Tooth formation occurs ontogenetically late in frogs, during or after metamorphosis, in contrast to during early larval or embryonic development in other vertebrates (Davit-Béal et al. 2007, Lainoff et al. 2015). This delayed shift in odontogenesis may be linked to the evolutionary lability of teeth in anurans. There may also be a relationship between the loss of teeth and delayed ossification of dentigerous elements. For example, the dentary bone ossifies relatively late in frogs, and nearly always lacks teeth, compared to being one of the first cranial elements to ossify in salamanders and caecilians (Harrington et al. 2013), and these amphibians always retain mandibular dentition. The anuran mouth undergoes dramatic restructuring during metamorphosis while transitioning from an herbivorous tadpole with a keratinized beak and short, cartilaginous lower jaw to a carnivorous frog with an elongated, bony lower jaw. This rapid morphological transformation requires further study in edentulous and toothed species. Several anuran lineages have evolved direct development (undergoing the larval stage within the egg; Gomez-Mestre et al. 2012), and this life history transition may provide an opportunity to repattern the jaw and alter dental development.

### Amphibian dentition and tooth loss in frogs

Dentition is highly conserved in salamanders and caecilians with no identified cases of edentulism. Teeth are present on the jaws and palate of all caecilians (Wake and Wurst 1979), and this is also the typical dental condition in salamanders (Gregory et al. 2016). The aquatic sirenid salamanders (*Siren, Pseudobranchus*) lack maxillary and premaxillary teeth (Clemen and Greven 1988), while the miniaturized species of *Thorius* sampled here (*T. pennatulus*) lacks maxillary teeth but retains teeth on the premaxilla, palate, and lower jaw. At least one species of *Thorius* possesses a novel dental polymorphism in which males lack maxillary teeth but females maintain several teeth on the maxilla (Hanken et al. 1999). To our knowledge, this is the only known case of a sexually dimorphic presence/absence dental polymorphism in an amphibian. Larval salamanders and caecilians were excluded in our dentition survey but differ in patterns of dentition from adults (Wake 1976, Clemen and Gren 2018), such as the transient presence of teeth on the splenials, palatines, and pterygoids that are lost during development (Schoch et al. 2019). Maxillary and premaxillary teeth are synchronized in all anuran taxa that we sampled, but two species in the genus *Telmatobius* have maxillary teeth in the absence of premaxillary teeth (Barrionuevo 2017). Members of this genus can be toothed or edentulous, and two species are reported to have intraspecific variation in the presence or absence of dentition (Barrionuevo 2017). Vomerine teeth are not coordinated with dentition on the upper jaw in frogs, are the most variable across the sampled anuran genera, and their lability requires further study. Previous work has suggested that the size and number of vomerine teeth may be correlated with diet and body size (Hedges 1989, Estrada and Hedges 1996). Teeth are entirely absent in 134 anuran genera distributed across 19 families, and our ancestral state reconstruction suggests that teeth have been lost more than 20 times during the evolution of frogs.

We identified a phylogenetic correlation between the evolution of edentulism and a microphagous diet, and these two traits co-occur in more than 50 genera belonging to 14 families (Dataset S2; Fig. 3). The majority of these species specialize on eating ants and termites, despite that these insects have many defense behaviors (biting, stinging, chemical weapons) and low nutritional value compared to other invertebrates (Redford and Dorea 1984, McNab 1984). Edentulous, microphagous frogs inhabit biomes ranging from tropical forests (e.g., *Dendrobates, Mantella, Cardioglossa*) to arid deserts (e.g., *Breviceps, Notaden*) and are found on all continents, excluding Antarctica. Frogs, ants, and termites evolved at roughly the same time—with important diversification events occurring in all three groups during the Cretaceous and Cretaceous–Paleogene boundary (Moreau and Bell 2013, Bourguignon et al. 2014, Feng et al. 2017)—suggesting the repeated evolution of complete edentulism in frogs may be linked to the spatiotemporal diversification of ants and termites. Teeth have been repeatedly reduced in other tetrapods that specialize on eating ants and termites, including multiple lineages of mammals (echidnas, numbats, aardvarks, aardwolves, anteaters, armadillos, pangolins; Reiss 2001) and squamates (scolecophidian blind snakes, *Aprasia* worm lizards; Daza and Bauer 2015).

The complete loss of teeth in frogs is associated with the shortening of the lower jaw (Fig. 4), a skeletal trait that is known to occur in species that eat smaller prey (Emerson 1985, Vidal-Garcia and Keogh 2017, Paluh et al. 2020). The shortening of the mandible reduces maximum gape and alters jaw biomechanics to improve the efficiency of catching many small prey items. Frogs with a jaw length equal to or longer than the skull have an asymmetrical feeding cycle where the time spent catching prey is short but the time spent bringing prey into the mouth is long (Gans and Gorniak 1982); shortened jaws result in a faster, symmetric feeding cycle where equal amounts of time are spent catching and bringing prey into the mouth (Emerson 1985). At least four lineages of edentulous anurans that specialize on ants and termites have additionally evolved muscular hydrostatic tongues that can be aimed in all three dimensions and with great precision without moving the head to improve the efficiency of small prey capture (*Rhinophrynus*, Trueb and Gans 1983; *Hemisus*, Nishikawa et al. 1999; microhylids and brevicipitids, Meyers et al. 2004). Frog species that feed inside ant and termite colonies may also possess improved abilities to process olfactory and tactical cues in order to detect and localize prey (Deban et al. 2001).

The majority of the 134 edentulous frogs in our dataset are restricted to the families Bufonidae and Microhylidae. All 48 genera of bufonids examined—the only anuran clade widely recognized as being edentulous (Davit-Béal et al. 2009)—and 48 of 54 of microhylid genera examined lack teeth. All remaining families have less than ten edentulous genera. The Bufonidae and Microhylidae are two of the most diverse frog families, comprising 638 and 695 species, respectively (18% of all frogs; AmphibiaWeb 2021). The evolution of edentulism in frogs may exert an influence on diversification rates, but we refrain from testing this hypothesis using trait-dependent diversification models due to our sparse, genus-level taxonomic sampling (429 tips representing 7,299 lineages). The results of our ancestral state reconstruction analyses indicate that teeth were independently lost in the most recent common ancestors of both bufonids and microhylids. Once lost, teeth have not been regained in the Bufonidae but may have re-evolved several times in microhylids. Although both clades have many taxa that specialize on microphagous prey, there are bufonids and microhylids with expanded, generalized diets, and a few species that will even consume vertebrate prey (e.g., *Rhinella marina, Asterophrys turpicola*). The variation in diet within bufonids and microhylids corresponds with variation in the relative length of the lower jaw (Dataset S3) and overall skull morphology (Paluh et al. 2020).

The inferred reversals in Microhylidae occur in *Dyscophus, Uperodon*, and four cophyline genera (*Anodonthyla, Cophyla, Plethodontohyla, Rhombophryne*). Recent work has also suggested that additional microhylid genera contain toothed and edentulous species (*Mini*, Scherz et al. 2019; *Glyphoglossus*, Gorin et al. 2021). If teeth were entirely lost in the common ancestor of microhylids, the repeated re-evolution of true teeth (with enamel, dentin, and pulp cavity) in this clade is unlikely and requires histological investigation. We hypothesize the tooth-like structures in these taxa may be small odontoid serrations, similar to what has been described in some *Brachycephalus* (Ribeiro et al. 2017) and New Guinea asterophrynine microhylids (Zweifel 1971). The dental anatomy of *Dyscophus* has been examined (LaDouceur et al. 2020), and this genus does possess true teeth. The phylogenetic relationships among microhylid taxa remain controversial (Peloso et al. 2016, Streicher et al. 2020), which further impedes the interpretation of dental evolution in this group.

Of the nine anuran genera known to possess variation in the presence or absence of teeth, diet data are only available for *Physalaemus* and *Engystomops*. In both genera, edentulous species have specialized microphagous diets in comparison to toothed congeners that consume a broader array of invertebrates (Narváez and Ron 2013). Dietary alkaloid sequestration has evolved as a predator defense mechanism in at least five clades of frogs that specialize on eating ants and mites, and teeth have been lost in several of these lineages (Dendrobatinae and *Ameerega* [Dendrobatidae, Saporito et al. 2004]; *Pseudophryne* [Myobatrachidae, Smith et al. 2002]; *Mantella* [Mantellidae, Daly et al. 1997]; *Melanophryniscus* [Bufonidae, Hantak et al. 2013]). Teeth are retained in *Phyllobates*, which is sister to all other edentulous genera of the Dendrobatinae, and in the Cuban *Eleutherodactylus* group known to sequester alkaloids (Rodríguez et al. 2010). There are other microphagous frogs that retain teeth, such as the ant specialist *Sphaenorhynchus* (Parmelee 1999). Further work is needed to investigate the number, size, and histological anatomy of teeth across toothed frogs that vary in diet. It remains unknown whether any anurans have lost enamel but retain teeth, which has occurred several times in mammals (Meredith et al. 2009).

No relationship was identified between complete edentulism and body size in the 423 frog species sampled. The smallest known species of frog, *Paedophryne amauensis*, lacks teeth, but some miniaturized anurans are toothed. We examined 25 taxa with a SVL of 15 mm or less: 13 were toothed and 12 were edentulous. Several of the smallest edentulous species in our dataset are microhylids, bufonids, and dendrobatids, and these clades have widespread tooth loss across a range of body sizes. We identified only one case of edentulism in Brachycephaloidea, in the genus *Brachycephalus*, despite that this new world radiation of over 1,000 species contains many miniaturized lineages (e.g., smallest members of *Pristimantis, Eleutherodactylus, Noblella*). Within the genus *Arthroleptis*, several miniature species lack teeth (Laurent 1954; Blackburn 2008), suggesting that, in some cases, a reduction in body size and tooth loss may be linked. There are several large or gigantic species within the Bufonidae (Womack and Bell 2020), but all true toads lack teeth regardless of size.

### Tooth loss in fossil frogs

Several crown-group fossil frogs have been described that are edentulous, including *Theatonius lancensis* (Fox 1976), *Tyrrellbatrachus brinkmani* (Gardner 2014), *Saltenia* (Baez 1981), and *Vulcanobatrachus* (Trueb et al. 2005) from the Late Cretaceous, and *Chelomophrynus bayi* from the Eocene (Henrici 1991). The majority of these taxa have been hypothesized to be members of the Pipoidea. Of the stem salientians with cranial material, teeth are present on the upper jaw in *Prosalirus* (Shubin and Jenkins 1995), *Vieraella* (Báez and Basso 1996), and *Liaobatrachus* (Gao and Wang 2001). No teeth are visible in *Triadobatrachus* (Ascarrunz et al. 2016), the oldest known stem frog, but the maxilla and premaxilla are poorly preserved in this impression fossil. The dentary of *Triadobatrachus* lacks teeth, and the absence of dentition on the lower jaw is considered a synapomorphy of Salientia (Milner 1988). To our knowledge, no stem tetrapods have been described as edentulous (Ruta et al. 2003, Anderson et al. 2008, Matsumoto and Evans 2017). Albanerpetontids, an extinct lineage of lissamphibians, retained teeth (Daza et al. 2020).

### Molecular and developmental mechanisms of tooth loss

Recent work has documented that several lineages of edentulous vertebrates have various states of molecular tooth decay in the genes that are critical for the formation of dentin and enamel with frameshift mutations and stop codons that result in nonfunctionalization (mammals: Meredith et al. 2009; turtles: Meredith et al. 2013; birds: Meredith et al. 2014; syngnathids: Lin et al. 2016). The frameshift mutation rate of these loci can be used to estimate the timing of tooth loss in the fossil record (Meredith et al. 2009, 2014), and the ratio of synonymous and nonsynonymous substitutions can be calculated to measure selection pressure on enamel matrix proteins (Alazem and Abramyan 2019). Whether edentulous frogs possess similar rates of molecular tooth decay in these loci, as demonstrated in amniotes, has yet to be tested. We hypothesize that these tooth-specific genes have degenerated repeatedly across edentulous anurans by novel inactivating mutations, and the frameshift mutation rate will indicate that teeth were lost at several different geologic times during the evolution of frogs. Anuran enamel matrix proteins may be operating under relaxed selection, compared to purifying selection in most mammals and reptiles (Alazem and Abramyan 2019), due to the evolution of projectile tongue feeding, enabling the evolutionary lability of frog teeth.

The developmental genetics of tooth formation in amphibians is almost entirely unexplored, especially when compared to our understanding of chondrichthyan, teleost, and amniote odontogenesis (Fraser et al. 2004, Tucker and Sharpe 2004, Thiery et al. 2017). It is unknown if the genes critical for tooth formation in fishes and amniotes are also expressed during morphogenesis of teeth in amphibians, if all frog species retain a suppressed ancestral developmental pathway of tooth development on the lower jaw, or if the odontogenetic pathway has been disrupted via one or many mechanisms on the jaws of edentulous anurans. The loss of teeth on the lower jaw of frogs could be due to the loss of a single major signal that can orchestrate odontogenesis, comparable to the sole loss of odontogenic *Bmp4* expression in living birds (Chen et al. 2007) or termination of *Msx2* expression in living turtles (Tokita et al. 2012), which arrests tooth formation early in development. If true, potential rudimentary structures, such as tooth buds or the early thickening of the odontogenic band, might be seen before the abortion of tooth development in the lower jaw of anurans. Investigation of the developmental genetics of tooth formation in the upper and lower jaws of frogs will fill a large gap in our understanding of vertebrate evolution and development and may elucidate the mechanisms of repeated tooth loss and putative cases of the re-evolution of lost teeth in one of the most diverse vertebrate orders.

## Methods

### Species Sampling and scanning

We collected data from high-resolution micro-computed tomography (microCT) scans of 523 amphibian species, representing 420 frog genera (of 461 total; AmphibiaWeb 2021), 65 salamander genera (of 68 total), and 30 caecilian genera (of 34 total). One recently described frog species was not CT scanned but included in the dataset because it is the only member of its genus with teeth (*Uperoleia mahonyi*; Clulow et al. 2016). All genera are represented by one species except for nine anuran genera (*Arthroleptis, Cacosternum, Engystomops, Gastrotheca, Physalaemus, Pipa, Telmatobius, Uperodon, Uperoleia*) with two sampled lineages that represent known dental variation within these genera (Dataset S1). All scans were run using a 240kv x-ray tube containing a diamond-tungsten target, with the voltage, current, and detector capture time adjusted for each scan to maximize absorption range for each specimen. Raw x-ray data were processed using GE’s proprietary datos|x software version 2.3 to produce a series of tomogram images and volumes, with final voxel resolutions ranging from 1 to 147 μm. The resulting microCT volume files were imported into VG StudioMax version 3.2.4 (Volume Graphics, Heidelberg, Germany), the skull and skeleton were isolated using VG StudioMax’s suite of segmentation tools, and then exported as high-fidelity mesh files. We deposited image stacks (TIFF) and 3D mesh files of the skull and skeleton for each specimen in MorphoSource (see Dataset S1 for DOIs).

### Survey of amphibian dentition variation and ancestral state reconstructions

We recorded the presence or absence of teeth on each dentigerous bone of the lower jaw, upper jaw, and palate for 524 amphibian species (Fig. 1; Dataset S1). We conducted ancestral state reconstructions of dentition (two states: toothed, edentulous) in extant amphibians using the data collected from 524 species representing 515 genera and all 77 families and the phylogeny of Jetz and Pyron (2018). Bayesian ancestral state reconstructions were calculated using reversible-jump MCMC in RevBayes (Höhna et al. 2016) to sample all five Markov models of phenotypic character evolution (one-rate, two-rate, zero-to-one irreversible, one-to-zero irreversible, no change) in proportion to their posterior probability. We accounted for model uncertainty by making model-averaged ancestral state estimates (Freyman and Höhna 2018, Freund et al. 2018). The models were assigned an equal prior probability using a uniform set-partitioning prior, and the root state frequencies were estimated using a flat Dirichlet prior. The rates of gain and loss of dentition were drawn from an exponential distribution with a mean of 10 expected character state transitions over the tree. The MCMC was run for 22,000 iterations, the first 2,000 iterations were discarded as burn-in, and samples were logged every 10 iterations. Convergence of the MCMC was confirmed using Tracer v1.6 to ensure that analyses had reached stationarity. We conducted additional ancestral state reconstructions to model the evolutionary history of dentition presence/absence on individual dentigerous elements (Figure 1—figure supplements 2–5).

### Testing relationships among edentulism, diet, and body size

We compiled dietary data for all sampled anuran species from the literature (see Dataset S2 for references). Species were classified as microphagous specialists if the majority (> 50%) of their diet by number or volume consists of ants, termites, or mites. Species were classified as generalists if the majority of their diet by number or volume consists of other invertebrate groups or vertebrates. For species with no published diet records, we searched for any existing diet records at the genus level. Due to the disparity in existing ecological data available across all anurans, the dietary records ranged from singular reports (one prey item in one individual) to detailed studies investigating the stomach contents of dozens of individuals through space and time.

We measured snout–vent length (SVL; tip of the snout to the rear of the ischium), skull length (occiput to tip of the snout) and mandible length (posterior to anterior tip of the lower jaw) for all sampled specimens using the linear measurement tools in VG StudioMax and MeshLab (Cignoni et al. 2008). We calculated relative jaw length (mandible length divided by skull length) for each specimen: a jaw length value greater than one indicates a posteriorly shifted jaw joint (lower jaw is longer than the head) and a value less than one indicates an anteriorly shifted jaw joint (lower jaw is shorter than the head).

We used phylogenetic comparative methods to test for evolutionary correlations among dentition, diet, and body size in frogs. We compiled diet records for 267 taxa, representing 258 genera and 52 anuran families: 158 species in the dentition dataset had published diet records and the remaining 109 lineages are represented by genus-level diet data. We excluded the remaining 162 anuran species in the dentition dataset (55 edentulous, 107 toothed) from the diet analyses due to the lack of known diet records at the species or genus level. Because dentition (toothed/edentulous) and diet (generalist/microphagous) were treated as binary traits, we tested for a phylogenetic correlation using discrete independent and discrete dependent models with rjMCMC sampling in BayesTraits v3.0.2 (Pagel and Meade 2006). The stepping stone sampler for marginal likelihood reconstructions was used with 100 stones and 1000 iterations. The branch lengths were scaled to have a mean of 0.1 using ScaleTrees. Bayes factors (Log BF = 2(log marginal likelihood complex model – log marginal likelihood simple model)) were used to compare the fit of the independent versus dependent models. Models were run using the complete 267 taxon dataset and a reduced 158 taxon dataset excluding genus-level diet data.

Several previous studies have demonstrated a correlation between skull shape and diet in frogs: species that specialize on small prey have anteriorly shifted, relatively short jaws while generalist feeders that are capable of eating large prey have a posteriorly shifted jaw joint (Emerson 1985, Vidal-Garcia and Keogh 2017, Paluh et al. 2020). Because diet data are lacking for many anuran genera, we additionally tested for a phylogenetic correlation between dentition and the relative length of the jaw as a morphological proxy for diet. Lastly, because teeth may be lost as a byproduct of miniaturization (Hanken and Wake 1993, Smirnov and Vasil’eva 1995), we tested for a phylogenetic correlation between dentition state and body size (SVL). Phylogenetic logistic regression models were calculated in the *phylolm* R package (Ho and Ané 2014) using dentition and measurement data for 423 anuran species. Dentition (toothed/edentulous) was treated as the binary response variable and the log transformed size metrics (relative jaw length, SVL) as continuous predictor variables. We used the “logistic_MPLE” method, which maximizes the penalized likelihood of the logistic regression, with a btol of 10, a log.alpha.bound of 10, and 1,000 bootstrap replicates.

## Acknowledgments

We thank all of the institutions, curators, and collection managers that loaned us specimens for this study. We thank Marta Vidal-Garcia for providing access to the *Spicospina flammocaerulea* scan. We thank the D.C.B. laboratory at the Florida Museum of Natural History for helpful comments that improved an earlier version of this manuscript.

## Data Availability Statement

Computed tomography data (tiff stacks and mesh files) have been deposited in MorphoSource (see Dataset 1 for DOIs). Data and scripts for all analyses are available on GitHub at https://github.com/dpaluh/edentulous_frogs.

## Supplemental Information

Dataset S1: Spreadsheet of specimens examined (524 amphibian species) with associated dentition data and MorphoSource DOIs.

Dataset S2: Compiled dietary data and references for 267 frog lineages.

Dataset S3: Measurement data (skull length, jaw length, SVL) for 423 frog species.

Datasets S1–3 and scripts for all analyses are available on GitHub at https://github.com/dpaluh/edentulous_frogs.

**Figure 1—figure supplement 1.**
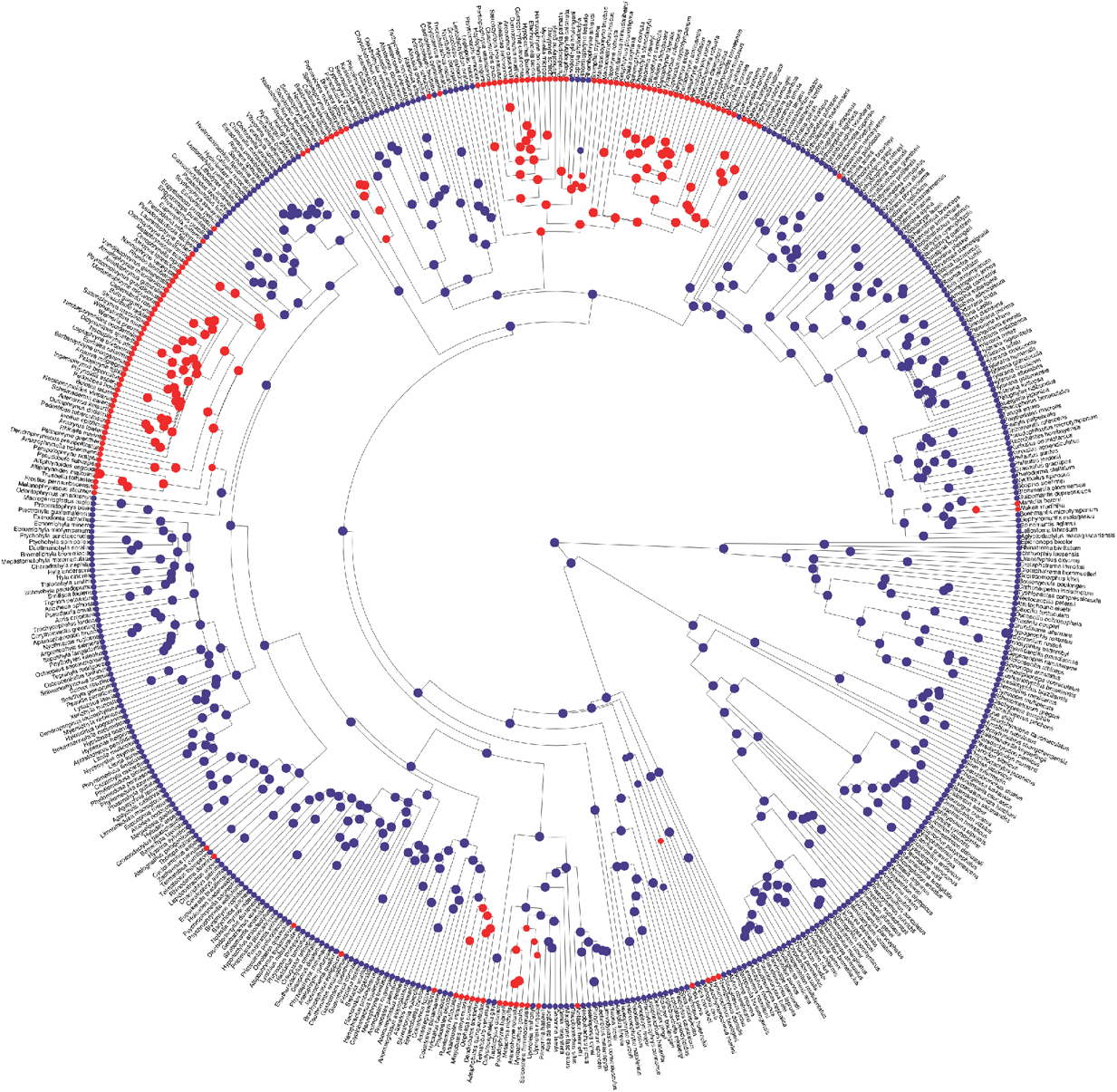
Phylogeny of 524 amphibians depicting the evolution of dentition (toothed = blue; edentulous = red) with species tip labels.

**Figure 1—figure supplement 2.**
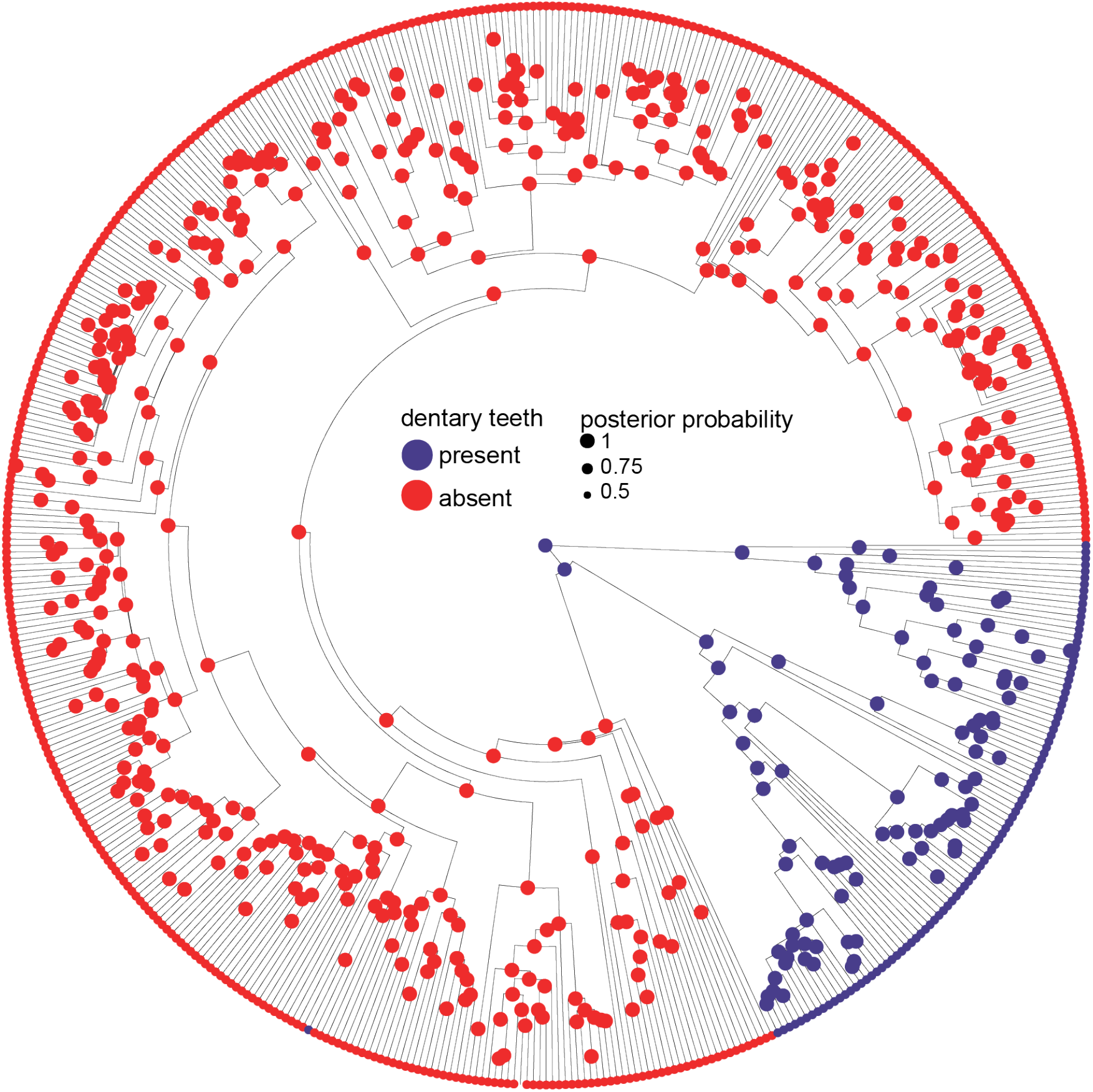
Phylogeny of 524 amphibians depicting the evolution of dentary teeth.

**Figure 1—figure supplement 3.**
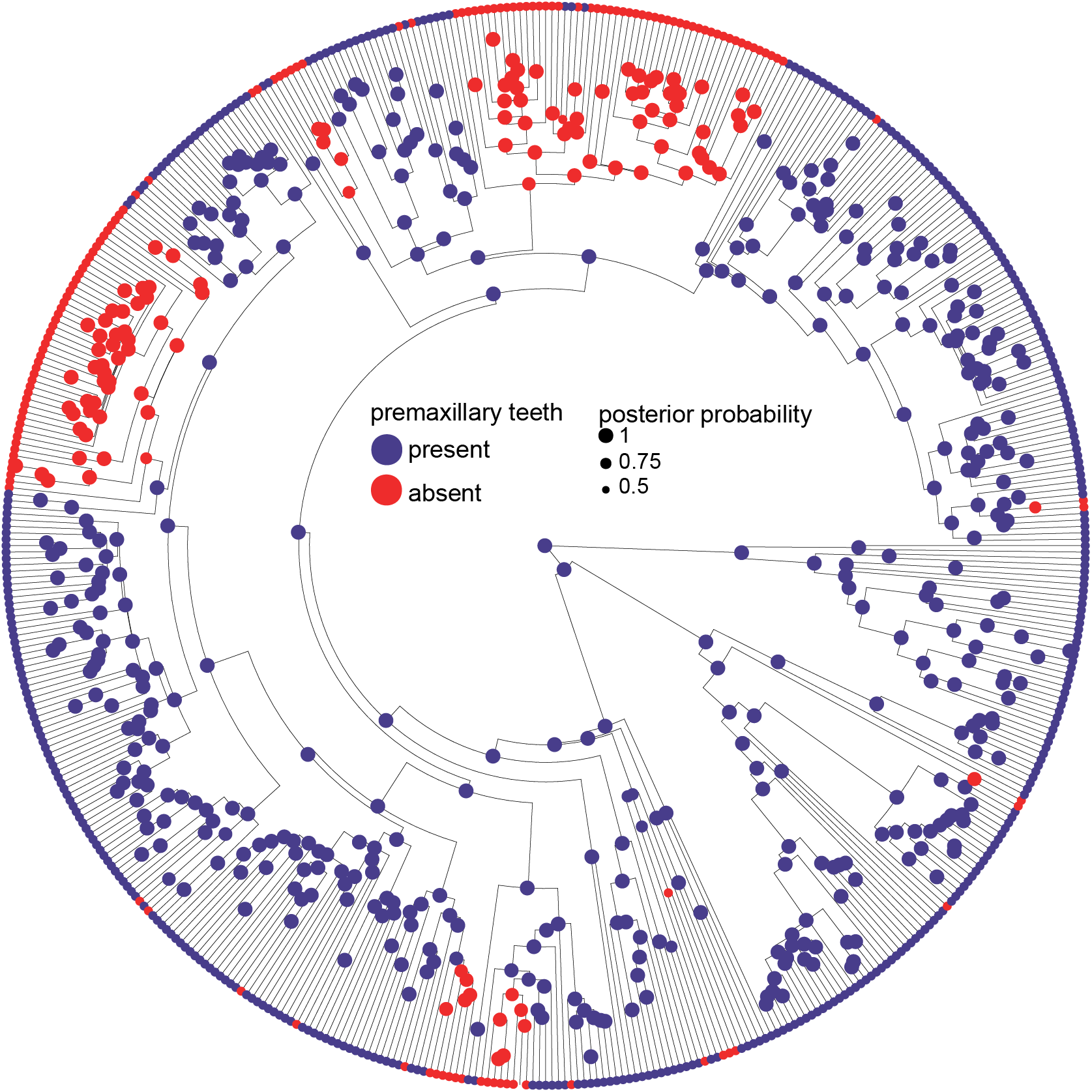
Phylogeny of 524 amphibians depicting the evolution of premaxillary teeth.

**Figure 1—figure supplement 4.**
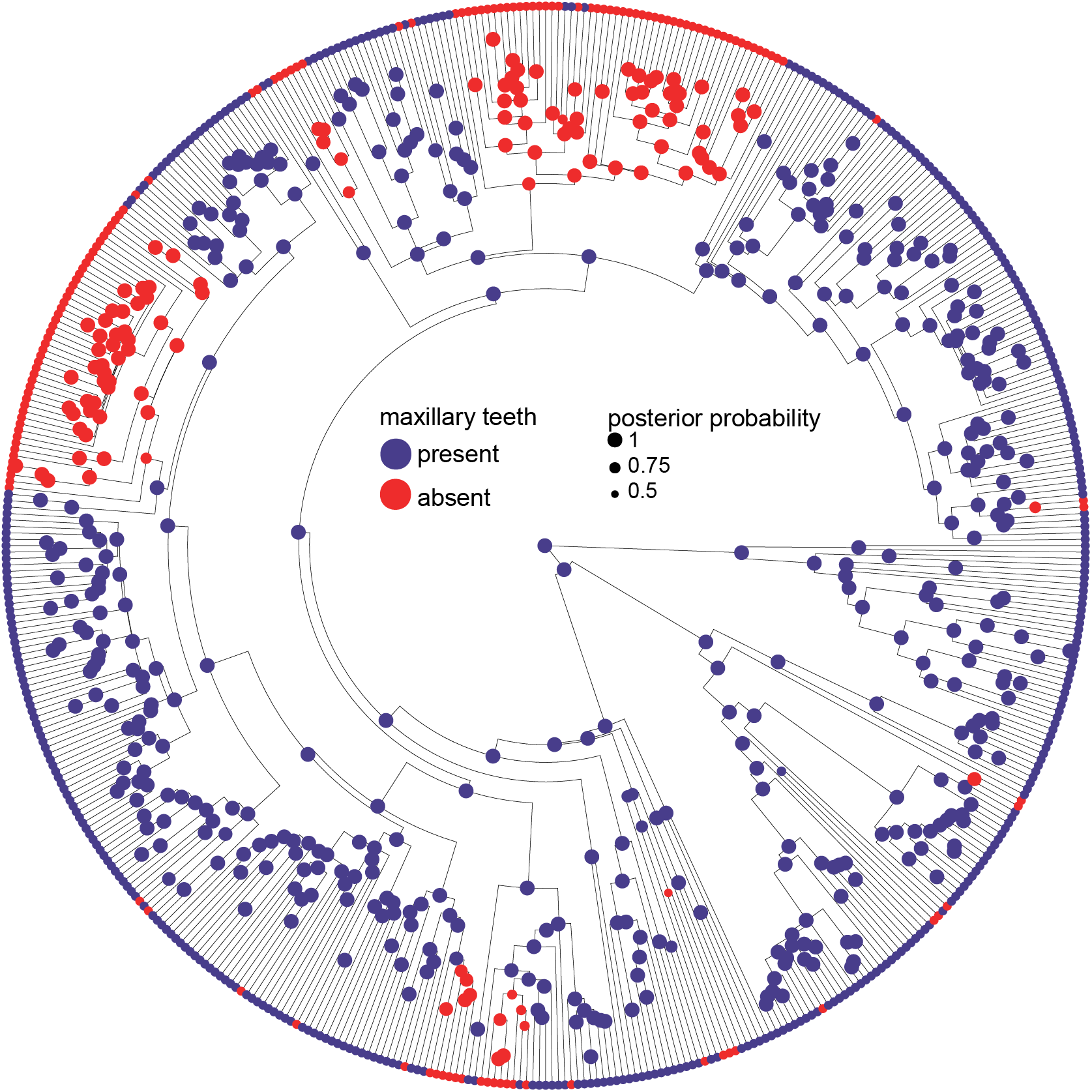
Phylogeny of 524 amphibians depicting the evolution of maxillary teeth.

**Figure 1—figure supplement 5.**
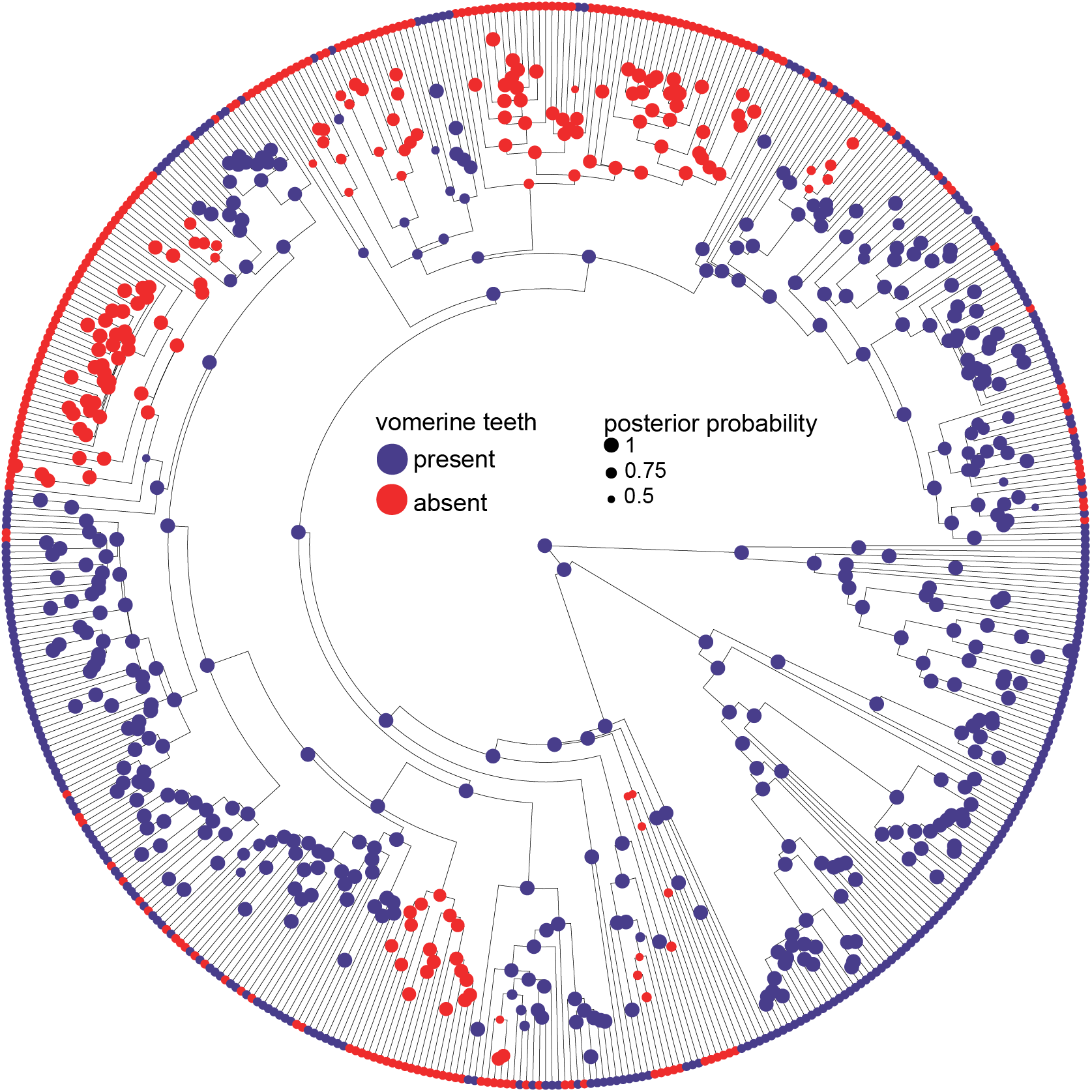
Phylogeny of 524 amphibians depicting the evolution of vomerine teeth.

**Figure 2—figure supplement 1.**
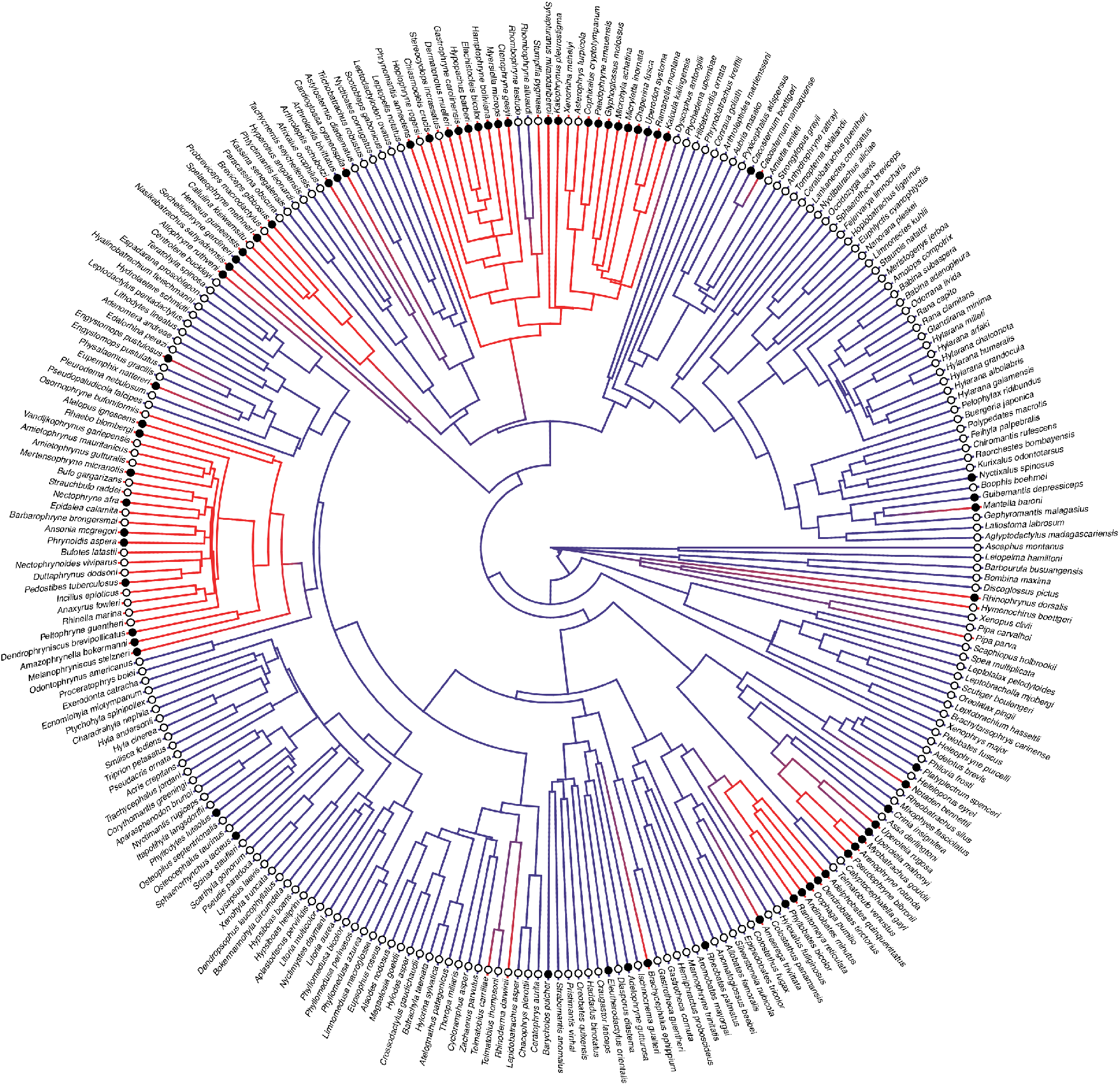
Phylogeny of 267 frog species with a stochastic character map of dentition and the distribution of diet states with species tip labels.

## References

Aigler, S.R., D. Jandzik, K. Hataa, K. Uesugi, D.W. Stock. 2014. Selection and constraint underlie irreversibility of tooth loss in cypriniform fishes. Proceedings of the National Academy of Sciences 111:7707–7712.

Alazem, O., J. Abramyan. 2019. Reptile enamel matrix proteins: selection, divergence, and functional constraint. Journal of Experimental Zoology B 2019:1–13.

Anderson, J.S., R.R. Reisz, D. Scott, N.B. Fröbisch, S.S. Sumida. 2008. A stem batrachian from the Early Permian of Texas and the origin of frogs and salamanders. Nature 453:515– 518.

Ascarrunz, E., J.C. Rage, P. Legreneur, M. Laurin. 2016. Triadobatrachus massinoti, the earliest known lissamphibian (Vertebrata: Tetrapoda) re-examined by μCT-Scan, and the evolution of trunk length in batrachians. Contributions to Zoology 85:201–234.

Báez, A.M. 1981. Redescription and relationships of Saltenia ibanezi, a Late Cretaceous pipid frog from northwestern Argentina. Ameghiniana 18:127–154.

Báez, A.M., N. Basso. 1996. The earliest known frogs of the Jurassic of South America: Review and cladistic appraisal of their relationships. Münchner geowissenschaftliche Abhandlungen A 30:131–158.

Blackburn, D.C. 2008. Evolution of diversity in African frogs (Arthroleptis and Cardioglossa). Ph.D. Dissertation, Harvard University, Cambridge, Massachusetts, 286 pp.

Barrionuevo, J.S. 2017. Frogs at the summits: phylogeny of the Andean frogs of the genus Telmatobius (Anura: Telmatobiidae) based on phenotypic characters. Cladistics 33:41– 68.

Boulenger, G.A. 1882. Catalogue of the Batrachia Salientia s. Ecaudata in the collection of the British Museum. 2nd ed. London: Taylor and Francis.

Bourguignon, T., N. Lo, S.L Cameron, J. Šobotník, Y. Hayashi, S. Shigenobu, D. Watanabe, Y. Roisin, T. Miura, T.A. Evans. 2014. The evolutionary history of termites as inferred from 66 mitochondrial genomes. Molecular Biology and Evolution 32:406–421.

Bhullar, B.-A. S., J. Marugán-Lobón, F. Racimo, G.S. Bever, T.B. Rowe, M.A. Norell, A. Abzhanov. 2012. Birds have paedomorphic dinosaur skulls. Nature 487:223–226.

Caldwell, J.P. 1996. The evolution of myrmecophagy and its correlates in poison frogs (family Dendrobatidae). Journal of Zoology 240:75–101.

Chen, Y., Y. Zhang, T.X. Jiang, A.J. Barlow, T.R. St. Amand, Y. Hu, S. Heaney, P. Francis-West, C.-M. Chuong, R. Maas. 2000. Conservation of early odontogenic signaling pathways in Aves. Proceedings of the National Academy of Sciences 97:10044–10049.

Cignoni, P., M. Callieri, M. Corsini, M. Dellepiane, F. Ganovelli, G. Ranzuglia. 2008. MeshLab: an Open-Source Mesh Processing Tool. Sixth Eurographics Italian Chapter Conference, 129–136.

Clemen, G.H. Greven, H. 1988. Morphological studies on the mouth cavity of Urodela. IX. Teeth of the palate and the splenials in Siren and Pseudobranchus (Sirenidae: Amphibia). Zeitschrift für zoologische Systematik und Evolutionsforschung 26:135–143.

Clemen, G., H. Greven. 2018. Long-term effects of arrested metamorphosis on dental systems in Salamandra salamandra (Salamandridae: Urodela). Vertebrate Zoology 68:143–155.

Clulow, S., M. Anstis, J.S. Keogh, R.A. Catullo. 2016. A new species of Australian frog (Myobatrachidae: Uperoleia) from the New South Wales mid-north coast sandplains. Zootaxa 4184:285–315.

Cope, E.D. 1867. On the families of the raniform Anura. Journal of the Academy of Natural Sciences of Philadelphia. Series 2. 6:189–206.

Cundall, D., E. Fernandez, F. Irish. 2017. The suction mechanism of the pipid frog, Pipa pipa (Linnaeus, 1758). Journal of Morphology 278:1229–1240.

Daly, J.W., H.M. Garraffo, G.S.E. Hall, J.F. Cover Jr. 1997. Absence of skin alkaloids in captive-raised Madagascan mantelline frogs (Mantella) and sequestration of dietary alkaloids. Toxicon 35:1131–1135.

Das, I., M. Coe. 1994. Dental morphology and diet in anuran amphibians from south India. Journal of Zoology London 233:417–427.

Davies, M. 1989. Ontogeny of bone and the role of heterochrony in the myobatrachine genera Uperoleia, Crinia, and Pseudophryne (Anura: Leptodactylidae: Myobatrachinae). Journal of Morphology 200:269–300.

Davit-Béal, T., H. Chisaka, S. Delgado, J.-Y. Sire. 2007. Amphibian teeth: current knowledge, unanswered questions, and some directions for future research. Biological Reviews 82:49–81.

Davit-Béal, T., AS Tucker, J-Y Sire. 2009. Loss of teeth and enamel in tetrapods: fossil record, genetic data and morphological adaptations. Journal of Anatomy 214:477–501.

Daza, J.D., A.M. Bauer. 2015. Cranial anatomy of the pygopodid lizard Aprasia repens, a gekkotan masquerading as a scolecophidian. In: O.R.P. Emonds, G.L. Powell, H.A. Jamniczky, A.M. Bauer, J. Theordor, eds. All animals are interesting: a Festschrift in honour of Anthony P. Russell. Oldenburg: BIS-Verlag der Carl von Ossietzky Universität, pp. 303–350.

Daza, J.D., E.L. Stanley, A. Bolet, A.M. Bauer, J.S. Arias, A. Čerňanský, J.J. Bevitt, P. Wagner, S.E. Evans. 2020. Enigmatic amphibians in mid-Cretaceous amber were chameleon-like ballistic feeders 370:587–691.

Dean, M.N. 2003. Suction feeding in the pipid frog, Hymenochirus boettgeri: Kinematic and behavioral considerations. Copeia 2003:879–886.

Deban, S.M., J.C. O’Reilly, K.C. Nishikawa. 2001. The evolution of the motor control of feeding in amphibians. American Zoologist 41:1280–1298.

Duellman, W.E., L. Trueb. 1986. Biology of Amphibians. New York: McGraw-Hill Book Co.

Emerson, S.B. 1985. Skull shape in frogs–correlations with diet. Herpetologica 1985:177–188.

Estrada, A.R., S.B. Hedges. 1996. At the lower size limit in tetrapods: A new diminutive frog from Cuba (Leptodactylidae: Eleutherodactylus). Copeia 1996:852–859.

Feng, Y-J., D.C. Blackburn, D. Liang, D.M. Hillis, D.B. Wake, D.C. Cannatella, P. Zhang. 2017. Phylogenomics reveal rapid, simultaneous diversification of three major clades of Gondwanan frogs at the Cretaceous-Paleogene boundary. Proceedings of the National Academy of Sciences 114:E5864–E5870.

Fordyce, R.E., L.G. Barnes. 1994. The evolutionary history of whales and dolphins. Annual Review of Earth and Planetary Sciences 22:419–455.

Fox, R.C. 1976. An edentulous frog (Theatonius lancensis, new genus and species) from the Upper Cretaceous Lance Formation of Wyoming. Canadian Journal of Earth Sciences 13:1486–1490

Fraser, G.J., A. Graham, M.M. Smith. 2004. Conserved deployment of genes during odontogenesis across osteichthyans. Proceedings of the Royal Society B 271:2311– 2317.

Freund, F.D., W.A. Freyman, C.J. Rothfels. 2018. Inferring the evolutionary reduction of corm lobation in Isoëtes using Bayesian model-averaged ancestral state reconstruction. American Journal of Botany 105:275–286.

Freyman, W.A., S. Höhna. 2018. Cladogenetic and anagenetic models of chromosome number evolution: A Bayesian model averaging approach. Systematic Biology 67:195–215.

Gans, C., G. Gorniak. 1982. Functional morphology of lingual protrusion in marine toads (Bufo mariunus). American Journal of Anatomy 163:195–222.

Gans, C., R.G. Northcutt. 1983. Neural crest and the origin of vertebrates: a new head. Science 220:268–273.

Gao, K.-Q., Y. Wang. 2001. Mesozoic anurans from Liaoning Province, China, and phylogenetic relationships of archaeobatrachian anuran clades. Journal of Vertebrate Paleontology 21:460–476.

Gardner, J.D. 2014. An edentulous frog (Lissamphibia; Anura) from the Upper Cretaceous (Campanian) Dinosaur Park Formation of southeastern Alberta, Canada. Canadian Journal of Earth Sciences 52:569–580.

Gomez-Mestre, I, R.A. Pyron, J.J. Wiens. 2012. Phylogenetic analyses reveal unexpected patterns in the evolution of reproductive modes in frogs. Evolution 66:3687–3700.

Gorin, V.A., M.D. Scherz, D.V. Korost, N.A. Poyarkov. 2021. Consequences of parallel minaturisation in Microhylinae (Anura, Microhylidae), with the description of a new genus of diminutive South East Asian frogs. Zoosystematics and Evolution 97:21–54.

Gregory, A.L., B.R. Sears, J.A. Wooten, C.D. Camp, A. Falk, K. O’Quin, T.K. Pauley. 2016. Evolution of dentition in salamanders: relative roles of phylogeny and diet. Biological Journal of the Linnean Society 119:960–973.

Hanken, J., D.B. Wake. 1993. Miniaturization of body size: organismal consequences and evolutionary significance. Annual Review of Ecology, Evolution, and Systematics 24:501–519.

Hanken, J., D.B. Wake, H.L. Freeman. 1999. Three new species of minute salamanders (Thorius: Plethodontidae) from Guerrero, México, including the report of a novel dental polymorphism in Urodeles. Copeia 1999:917–931.

Hantak, M.M., T. Grant, S. Reinsch, D. McGinnity, M. Loring, N. Toyooka, R.A. Saporito. 2013. Dietary alkaloid sequestration in a poison frog: An experimental test of alkaloid uptake in Melanophryniscus stelzneri (Bufonidae). Journal of Chemical Ecology 39:1400–1406.

Harrington, S.M., L.B. Harrison, C.A. Sheil. 2013. Ossification sequence heterochrony among amphibians. Evolution and Development 15:344–364.

Hedges, S.B. 1989. Evolution and biogeography of West Indian frogs of the genus Eleutherodactylus: slow-evolving loci and the major groups. In C.A. Woods, ed., Biogeography of the West Indies: past present and future. Gainesville, Florida: Sandhill Crane Press, pp. 305–370.

Hendrickx, C., O. Mateus, R. Araújo, J. Choiniere. 2019. The distribution of dental features in non-avian theropod dinosaurs: Taxonomic potential, degree of homoplasy, and major evolutionary trends. Palaeontologia Electronica 22:1–110. doi:10.26879/820.

Henrici, A.C. 1991. Chelomophrynus bayi (Amphibia, Anura, Rhinophrynidae), a new genus and species from the middle Eocene of Wyoming: ontogeny and relationships. Annals of the Carnegie Museum 60:97–144.

Hime, P.M., A.R. Lemmon, L.E.C. Moriarty, E. Prendini, J.M. Brown, R.C. Thomson, J.D. Kratovil, B.P. Noonan, R.A. Pyron, P.L.V. Peloso, M.L. Kortyna, J.S. Keogh, S.C. Donnellan, M.R. Lockridge, C.J. Raxworthy, K. Kunte, S.R. Ron, S. Das, N. Gaitonde, D.M. Green, J. Labisko, J. Che, D.W. Weisrock. Phylogenomics reveals ancient gene tree discordance in the amphibian tree of life. Systematic Biology 70:49–66.

Ho, L.S.T., C. Ané. 2014. A linear-time algorithm for Gaussian and non-Gaussian trait evolution models. Systematic Biology 63:397–408.

Höhna, S., M.J. Landis, T.A. Heath, B. Boussau, N. Lartillot, B.R. Moore, J.P. Huelsenbeck, F. Ronquist. 2016. RevBayes: Bayesian phylogenetic inference using graphical models and an interactive model-specification language. Systematic Biology 65:726–736.

Jetz, W., R.A. Pyron. 2018. The interplay of past diversification and evolutionary isolation with present imperilment across the amphibian tree of life. Nature Ecology and Evolution 2:850–858.

Kohno, H., R. Ordonio-Aguilar, A. Ohno, Y. Taki. Morphological aspects of feeding and improvement in feeding ability in early stage larvae of the milkfish, Chanos chanos. Ichthyological Research 43:133–140.

Kottelat, M., R. Britz, T.H. Hui, K.-E. Witte. 2006. Paedocypris, a new genus of Southeast Asian cyprinid fish with a remarkable sexual dimorphism, comprises the world’s smallest vertebrate. Proceedings of the Royal Society B 273:895–899.

LaDouceur, E.E.B., A.M. Hauck, M.M. Garner, A.N. Cartoceti, B.G. Murphy. 2020. Odontomas in Frogs. Veterinary Pathology 57:147–150.

Lainoff, A.J., J.E. Moustakas-Verho, D. Hu, A. Kallonen, R.S. Marcucio, L.J. Hlusko. 2015. A comparative examination of odontogenic gene expression in both toothed and toothless amniotes. Journal of Experimental Zoology B 324:255–269.

Laurent, R.F. 1954. Remarques sur le genre Schoutedenella Witte. Annales du Musée Royal du Congo Belge, 4, Sciences Zoologiques, Tervuren 1:34–40.

Lawson, R., D.B. Wake, N.T. Beck. 1971. Tooth replacement in the Red-backed Salamander, Plethodon cinereus. Journal of Morphology 134:259–270.

Lin, Q., S. Fan, Y. Zhang, M. Xu, H. Zhang, Y. Yang, et al. 2016. The seahorse genome and the evolution of its specialized morphology. Nature 540:395–399.

Matsumoto, R., S.E. Evans. 2017. The palatal dentition of tetrapods and its functional signi?cance. Journal of Anatomy 230:47–65.

McNab, B.K. 1984. Physiological convergence amongst ant-eating and termite-eating mammals. Journal of Zoology 203:485–510.

Mihalitsis, M. D. Bellwood. 2019. Functional implications of dentition-based morphotypes in piscivorous fishes. Royal Society Open Science 6:190040.

Milner, A.R. 1988. The relationships and origin of living amphibians. In M.J. Benton, ed., The phylogeny and classification of the Tetrapods. Oxford: Clarendon Press, pp 59–102.

Meredith, R.W., J. Gatesy, W.J. Murphy, O.A. Ryder, M.S. Springer. 2009. Molecular decay of the tooth gene enamelin (ENAM) mirrors the loss of enamel in the fossil record of placental mammals. Plos Genetics 5:e1000634.

Meredith, R.W., J. Gatesy, M.S. Springer. 2013. Molecular decay of enamel matrix protein genes in turtles and other edentulous amniotes. BMC Evolutionary Biology 13:20.

Meredith, R.W., G. Zhang, M.T.P. Gilbert, E.D. Jarvis, M.S. Springer. 2014. Evidence for a single loss of mineralized teeth in the common avian ancestor. Science 346:1254390.

Meyers, J.J., J.C. O’Reilly, J.A. Monroy, K.C. Nishikawa. 2004. Mechanism of tongue protraction in microhylid frogs. Journal of Experimental Biology 207:21–31.

Moreau, C.S., C.D. Bell. 2013. Testing the museum versus cradle biological diversity hypothesis: Phylogeny, diversification, and ancestral biogeographic range evolution of the ants. Evolution 67:2240–2257.

Mulas, A., A. Bellodi, C. Porcu, A. Cau, E. Coluccia, R. Demurtas, M.F. Marongiu, P. Pesci, M.C. Follesa. 2020. Living naked: first case of lack of skin-related structures in an elasmobranch, the blackmouth catshark (Galeus melastomus). Journal of Fish Biology 97:1252–1256.

Narváez, A.E., S.R. Ron. 2013. Feeding habits of Engystomops pustulatus (Anura: Leptodactylidae) in western Ecuador. South American Journal of Herpetology 8:161– 167.

Nesbitt, S., M.A. Norell. 2006. Extreme convergence in the body plans of an early suchian (Archosauria) and ornithomimid dinosaurs (Theropoda). Proceedings of the Royal Society B 273:1045–1048.

Nishikawa, K.C., W.M. Kier, K.K. Smith. 1999. Morphology and mechanics of tongue movement in the African pig-nosed frog Hemisus marmoratum: a muscular hydrostatic model. Journal of Experimental Biology 202:771–780

Pagel, M., A. Meade. 2006. Bayesian analysis of correlated evolution of discrete characters by reversible-jump Markov chain Monte Carlo. The American Naturalist 167:808–825.

Paluh, D.P., E.L. Stanley, D.C. Blackburn. 2020. Evolution of hyperossification expands skull diversity in frogs. Proceedings of the National Academy of Sciences 117:8554–8562.

Parmelee, J.R. 1999. Trophic ecology of a tropical anuran assemblage. Scientific Papers, Natural History Museum, The University of Kansas 11:1–59.

Peloso, P.L.V., D.R. Frost, S.J. Richards, M.T. Rodrigues, S. Donnellan, M. Matsui, et al. 2016. The impact of anchored phylogenomics and taxon sampling on phylogenetic inference in narrow-mouthed frogs (Anura, Microhylidae). Cladistics. 32:113–140

Rauhut, O.W.M., A.M. Heyng, A. López-Arbarello, A. Hecker. 2012. New Rhynchocephalian from the Late Jurassic of Germany with a Dentition That Is Unique amongst Tetrapods. Plos One 7:e46839.

Redford, K.H., J.G. Dorea. 1984. The nutritional value of invertebrates with emphasis on ants and termites as food for mammals. Journal of Zoology 203:385–395.

Regal, P.J., C. Gans. 1976. Functional aspects of the evolution of frog tongues. Evolution 30:718–734.

Reiss, K.Z. 2001. Using phylogenies to study convergence: the case of the ant-eating mammals. American Zoologist 41:507–525.

Revell, L.J. 2012. phytools: An R package for phylogenetic comparative biology (and other things). Methods in Ecology and Evolution 3:217–223.

Ribeiro, L.F., D.C. Blackburn, E.L. Stanley, M.R. Pie, M.R. Bornschein. 2017. Two new species of the Brachycephalus pernix group (Anura: Brachycephalidae) from the state of Paraná, southern Brazil. PeerJ 5:e3603.

Rittmeyer, E.N., A. Allison, M.C. Gründler, D.K. Thompson, C.C. Austin. 2012. Ecological guild evolution and the discovery of the world’s smallest vertebrate. Plos One 7:e29797.

Rodríguez, A., D. Poth, S. Schulz, M. Vences. 2010. Discovery of skin alkaloids in a miniaturized eleutherodactylid frog from Cuba. Biology Letters 7:414–418.

Roos, G., S.V. Wassenbergh, A. Herrel, P. Aerts. 2009. Kinematics of suction feeding in the seahorse Hippocampus reidi. The Journal of Experimental Biology 212:3490–3498.

Rücklin, M., P.C.J. Donoghue, Z. Johanson, K. Trinajstic, F. Marone, M. Stampanoni. 2012. Development of teeth and jaws in the earliest jawed vertebrates. Nature 491:748–751.

Ruta, M., J.E. Jeffery, M.I. Coates. 2003. A supertree of early tetrapods. Proceedings of the Royal Society B 270:2507–2516.

Saporito, R.A., H.M. Garraffo, M.A. Donnelly, A.L. Edwards, J.T. Longino, J.W. Daly. 2004. Formicine ants: An arthropod source for the pumiliotoxin alkaloids of dendrobatid poison frogs. Proceedings of the National Academy of Sciences 101:8045–8050.

Scherz, M.D., C.R. Hutter, A. Rakotoarison, J.C. Riemann, M.-O. Rödel, S.H. Ndriantsoa, J. Glos, S.H. Roberts, A. Crottini, M. Vences, F. Glaw. 2019. Morphological and ecological convergence at the lower size limit for vertebrates highlighted by five new miniaturised microhylid frog species from three different Madagascan genera. Plos One 14:e0213314.

Schoch, R.R., P. Pogoda, A. Kupfer. 2019. The impact of metamorphosis on the cranial osteology of giant salamanders of the genus Dicamptodon. Acta Zoologica 2021:88– 104.

Shubin, N.H. F.A. Jenkins Jr. 1995. An Early Jurassic jumping frog. Nature 377:49–52.

Smith, B.P., M.J. Tyler, T. Kaneko, H.M. Garraffo, T.F. Spande, J.W. Daly. 2002. Evidence for biosynthesis of pseudophrynamine alkaloids by an Australian myobatrachid frog (Pseudophryne) and for sequestration of dietary pumiliotoxins. Journal of Natural Products 65:439–447.

Streicher, J.W., S.P. Loader, A. Varela-Jaramillo, P. Montoya, R.O. de Sá. 2020. Analysis of ultraconserved elements supports African origins of narrow-mouthed frogs. Molecular Phylogenetics and Evolution 146:106771.

Smirnov, S.V., A.B. Vasil’eva. 1995. Anuran dentition: development and evolution. Russian Journal of Herpetology 2:120–128.

Thiery, A.P., T. Shono, D. Kurokawa, R. Britz, Z. Johanson, G.J. Fraser. 2017. Spatially restricted dental regeneration drives pufferfish beak development. Proceedings of the National Academy of Sciences 114:4425–4434.

Tokita, M., W. Chaeychomsri, J. Siruntawineti. 2012. Developmental basis of toothlessness in turtles: insight into convergent evolution of vertebrate morphology. Evolution 67:260– 273.

Trueb, L., C. Gans. 1983. Feeding specializations of the Mexican burrowing toad, Rhinophrinus dorsalis (Anura: Rhinophrynidae). Journal of Zoology 199:189–208.

Trueb, L., C.F. Ross, R.M.H. Smith. 2005. A new pipoid anuran from the Late Cretaceous of South Africa. Journal of Vertebrate Paleontology 25:533–547.

Tucker, A., P. Sharpe. 2004. The cutting-edge of mammalian development; how the embryo makes teeth. Nature Reviews Genetics 5:499–508.

University of California, Berkeley. 2021. AmphibiaWeb: Information on amphibian biology and conservation. Available at amphibiaweb.org. Accessed January 15, 2021.

Vidal-Garcia, M., J.S. Keogh. 2017. Phylogenetic conservatism in skulls and evolutionary lability in limbs – morphological evolution across an ancient frog radiation is shaped by diet, locomotion and burrowing. BMC Evolutionary Biology 17:165.

Vences, M., F. Glaw,W. Böhme. 1998. Evolutionary correlates of microphagy in alkaloid-containing frogs (Amphibia: Anura). Zoologischer Anzeiger 236:217–230.

Visser, J. 1981. Tooth counts for Dasypeltis (Serpentes: Dasypeltinae). The Journal of the Herpetological Association of Africa 25:13–14.

Wake, M.H. 1976. The development and replacement of teeth in viviparous caecilians. Journal of Morphology 148:33–63.

Wake, M.H., G.Z. Wurst. 1979. Tooth crown morphology in caecilians (Amphibia: Gymnophiona). Journal of Morphology 159:331–341.

Wang, S., J. Stiegler, P. Wu, C.-M. Chuong, D. Hu, A. Balanoff, Y. Zhou, X. Xu. 2017. Heterochronic truncation of odontogenesis in theropod dinosaurs provides insight into the macroevolution of avian beaks. Proceedings of the National Academy of Sciences 114:10930–10935.

Wiens, J.J. 2011. Re-evolution of lost mandibular teeth in frogs after more than 200 million years, and re-evaluating Dollo’s Law. Evolution 65:1283–1296.

Womack, M.C., R.C. Bell. 2020. Two-hundred million years of anuran body-size evolution in relation to geography, ecology and life history. Journal of Evolutionary Biology 33:1417– 1432.

Yang, T.-R., P.M. Sander. 2018. The origin of the bird’s beak: new insights from dinosaur incubation periods. Biology Letters 14:20180090.

Zweifel, R.G. 1971. Relationships and distribution of Genyophryne thomsoni, a microhylid frog of New Guinea. American Museum Novitates 2469:1–13.

